# Coarse-grained implicit solvent lipid force field with a compatible resolution to the Cα protein representation

**DOI:** 10.1101/2020.09.20.305060

**Authors:** Diego Ugarte La Torre, Shoji Takada

## Abstract

Biological membranes have been prominent targets for coarse-grained (CG) molecular dynamics (MD) simulations. While minimal CG lipid models with three-beads per lipid and quantitative CG lipid models with >10-beads per lipid have been well studied, in between them, CG lipid models with a compatible resolution to residue-level CG protein models are much less developed. Here, we extended a previously developed three-bead lipid model into a five-bead model and parametrized it for two phospholipids, POPC and DPPC. The developed model, iSoLF, reproduced the area per lipid, hydrophobic thickness, and phase behaviors of the target phospholipid bilayer membranes at the physiological temperature. The model POPC and DPPC membranes were in liquid and gel phases, respectively, in accordance with experiments. We further examined the spontaneous formation of a membrane bilayer, the temperature dependence of physical properties, vesicle dynamics, and the POPC/DPPC two-component membrane dynamics of the CG lipid model, showing some promise. Once combined with standard Cα protein models, the iSoLF model will be a powerful tool to simulate large biological membrane systems made of lipids and proteins.

## Introduction

While molecular dynamics (MD) simulations have been indispensable tools to study the structural dynamics of biomolecular systems, the time scale attainable by all-atom MD simulations poses one of the major challenges for many biological phenomena.^1-3^ To overcome this limitation, coarse-grained (CG) modeling reduces the number of degrees of freedom by grouping atoms into CG beads, effectively decreasing the computational cost of simulations while retaining the properties of interest as much as possible.^5-6^ Due to the hierarchic nature of biomolecular systems, there are several different resolutions of coarse-graining. In general, higher resolution CG models are more accurate but computationally more expensive. Thus, depending on the purpose, one can choose the best CG model, among others. For example, CG models that explicitly represent solvent molecules are relatively accurate, while implicit solvent CG models are considerably faster by incorporating the average effects of solvents directly into CG force fields of solute molecules.^7^

Biological membranes are prominent targets of CG MD simulations, for which different classes of CG lipid models have been developed for two decades. In 1998, Goetz and Lipowsky developed an explicit solvent CG amphiphile model and successfully simulated self-assembly of a bilayer membrane.^8^ Some years later, Noguchi and Takasu were the first to make an implicit solvent CG model of amphiphiles that exhibit proper physical behaviors of a bilayer membrane.^9^ Later, Cooke et al. developed a much simpler pairwise-interacting implicit solvent CG model for lipids.^10^ Both of these implicit solvent models use three CG beads per lipid, making them minimal and generic, without requiring the parameterization for any specific molecule. These minimal models were successfully applied to uncover many physical aspects of membrane systems, such as the gel-liquid phase transition, phase separation, membrane fusion, and budding. As a different class of models, several higher-resolution CG lipid models were developed, including the seminal work of MARTINI by Marrink et al. in 2004.^11-18^ This class of models uses more than 10 CG beads per lipid and represent the two-alkyl-tail geometry explicitly, making the model specific to individual phospholipids. Among others, the MARTINI model has been successfully applied to many targets.^19-22^ Most of these models, with some exceptions^13,17,23^, use explicit solvent molecules, making them computationally demanding compared to the above-mentioned minimal models.

Notably, most biological membrane systems of interest contain membrane proteins as well. Thus, to be able to apply CG lipid models to many of these biological systems, its compatibility with CG protein models is of crucial importance. In particular, to model physicochemical interactions between lipids and proteins naturally, it is highly desired that both the CG lipid and protein representations share a similar resolution. The MARTINI force field, for example, consistently uses a mapping of one CG particle for about four non-hydrogen atoms for lipids, proteins, and other molecules. Among many CG models for proteins, a classic and still very popular representation is to use one CG particle per amino acid, most frequently placing the CG particle at its Cα position^24-28^. Amino acids in proteins contain 8.4 ± 2.4 non-hydrogen atoms (the average over 20 amino acids ± the standard deviation). Therefore, with this resolution, one can roughly reduce the degree of freedom by one order of magnitude. Representative phospholipids, for example, POPC (1-palmitoyl-2-oleoyl-sn-glycero-3-phosphocholine) and DPPC (1,2-dipalmitoyl-sn-glycero-3-phosphatidylcholine), contain 52 and 50 non-hydrogen atoms, respectively. We thus regard the use of 5-6 CG particles for each lipid molecule to be compatible with the Cα protein representation. However, among the several CG lipid models with an intermediate resolution in between three-beads per lipid and MARTINI-like models^29-35^, there are few CG models with a 5-6 CG particles per lipid resolution. In fact, the purpose of this paper is to present a new and relatively simple CG lipid force field with a compatible resolution to Cα protein models, where we represent each lipid with five CG beads to reduce the computational cost.

With the use of five CG beads, our aim is not to model generic lipid molecules but to parametrize the model for specific phospholipid molecules. In particular, we parametrize our CG lipid model for the POPC and the DPPC lipids. POPC and DPPC are unsaturated and saturated phospholipids, respectively. It is well-known that near-physiological temperatures (30 °C for example), pure POPC lipid membranes are in the liquid disordered phase, while pure DPPC lipid membranes are in the gel phase.^36,37^ More generally, at physiological conditions, pure unsaturated phospholipid membranes are in the liquid disordered phase, while pure saturated phospholipid membranes are in the gel phase. Reproducing these two phases should be important for simulations of biological membranes, which are a mixture of unsaturated and saturated phospholipids, in addition to membrane proteins and others. We also note that, with a five CG bead representation, we give up the two-tailed branched-chain geometry and, instead, use a linear chain (notably, it is not impossible to take two-tailed chain geometry with six beads per lipid resolution, as was recently proposed in an elegant work^34^). The linear chain representation of lipids makes it particularly challenging to distinguish between the unsaturated and saturated lipids because the unsaturated tail tends to bend and separate from the other tail.

In this work, we develop our CG implicit solvent lipid model by extending the work of Cooke, Kremer, and Deserno^10^, representing each lipid molecule with five CG beads so that it has a compatible resolution to Cα protein models. The Method section describes the CG model, including its mapping to all-atom structures and the potential energy function, as well as CG and all-atom MD simulation details. The Results and Discussion section begins with the parametrization of the force field for the two target lipids, POPC and DPPC. Then we report simulation results of spontaneous bilayer membrane formation, 2D diffusion of lipids, vesicle dynamics, the temperature dependence of membrane properties, and the POPC/DPPC two-component membrane dynamics. Finally, we discuss the limitation and future directions.

## Methods

### Lipid model

In our CG model, a two-tailed glycerophospholipid is represented as a linear chain molecule (**Fig. 1a**). Each CG lipid molecule is composed of five beads, two polar head beads (H1 and H2), and three hydrophobic tail beads (T1, T2, and T3). The H1 bead represents the terminal group bonded to the phosphate, and the H2 bead corresponds to the phosphate, glycerol, and ester carbonyls. The T1, T2, and T3 beads represent the first five carbon atoms of each tail, the next five carbon atoms of each tail, and the remaining carbon atoms, respectively. As mentioned, this five-bead mapping produces a similar resolution to Cα protein models.

**Figure 1.**
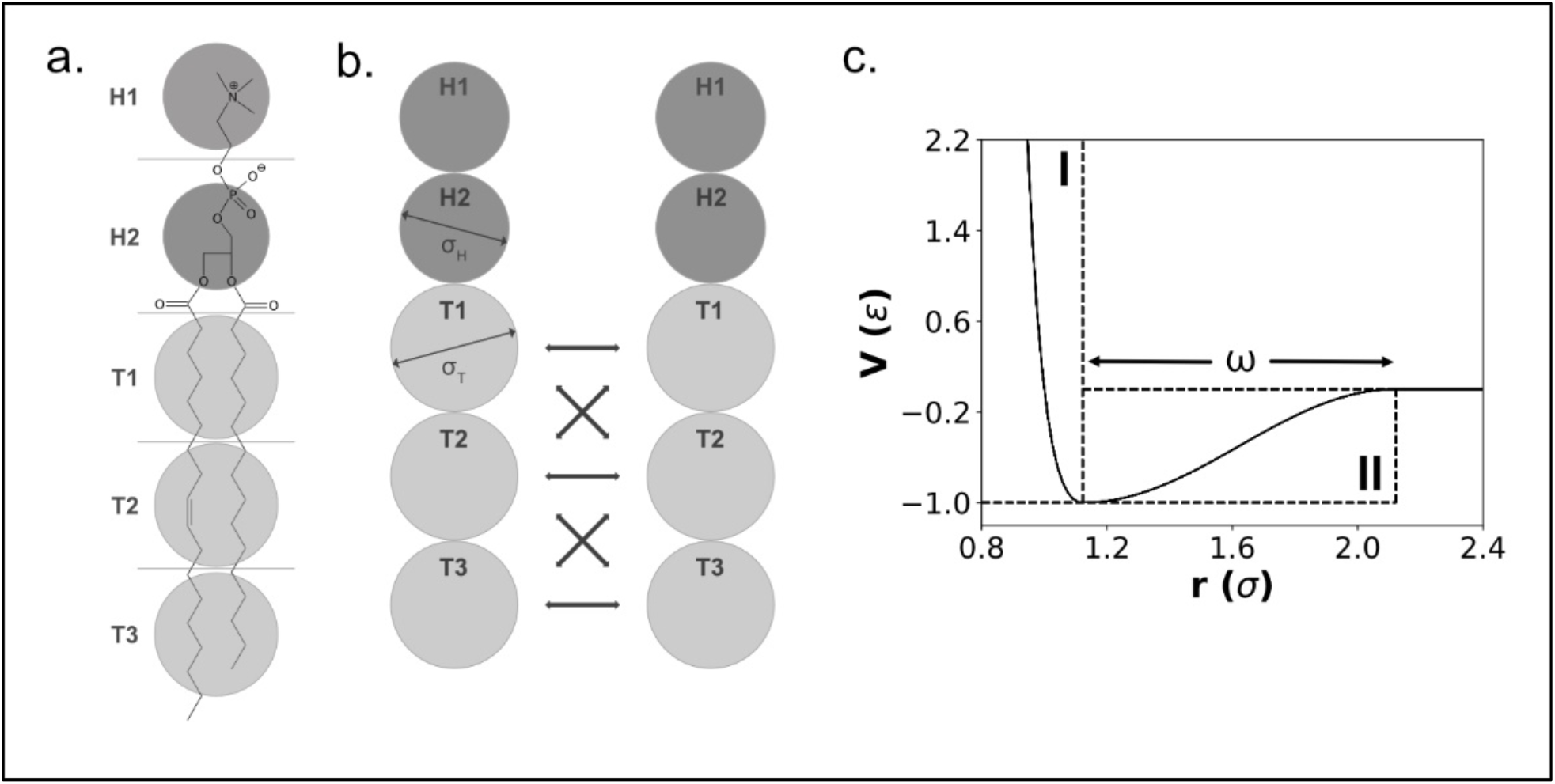
The current CG lipid model. (a) Mapping of glycerophospholipids, POPC as an example, into a linear chain of five CG beads composed of two polar head beads (H1 and H2), and three hydrophobic tail beads (T1, T2, and T3). The horizontal lines indicate the boundaries that define the assignment of atoms into CG beads. (b) Schematic picture of intermolecular interactions. The thick double-headed arrows indicate attractive interactions between hydrophobic tail CG particles. Notably, no attraction is applied between the T1 and T3 beads. Repulsion between any pairs is applied based on the bead diameter, *σ*_*H*_ for the head, and *σ*_*T*_, for the tail. (c) The potential function between tail CG beads (except T1-T3 pairs) in the unit of the scaling parameters *σ* and ε^10^. The regions **I** and **II** represent the repulsive and attractive parts, respectively. The width of the attractive part is controlled by the parameter ω.

In our CG force field, the potential energy function has four terms:

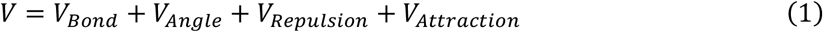

The first term, *V*_*Bond*_, represents the virtual bond interactions between two adjacent CG beads belonging to the same lipid molecule and is modeled by the harmonic potential:

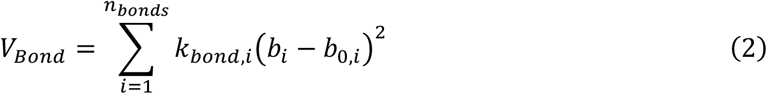

Here, *k*_*bond,i*_ is the force constant, *b*_*i*_ is the *i*-th virtual bond length between consecutive CG beads, *b*_0,*i*_ is the equilibrium value for the virtual bond, and *n*_*bonds*_ is the total number of virtual bonds. The second term, *V*_*Angle*_, is the potential for the virtual bond-angle between two consecutive virtual bonds in a lipid molecule and is modeled by the harmonic potential:

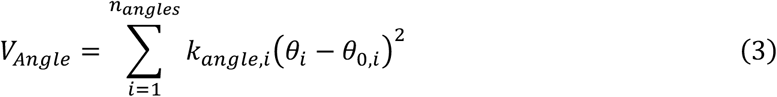

Here, *k*_*angle,i*_ is the force constant, *θ*_*i*_ is the angle between two consecutive bonds, *θ*_0,*i*_ is the equilibrium value for the *i*-th angle, and *n*_*angles*_ is the total number of angles. We note that the dihedral-angle potential used in all-atom force fields is not included in our model because the *θ*_0,*i*_ in eq.(3) turn out to be close to π (see Table 1, below), which makes the dihedral potential near divergent.

**Table 1.** Parameters for the virtual bond and bond-angle potentials of POPC and DPPC. The force coefficients *k* are in *kcal*/Å^2^*mol* for the virtual bond, and in *kcal*/*mol* for the virtual bond-angle, the equilibrium distances *b*_0_ are in Å, and the equilibrium angles *θ*_0_ are in radians.

| Type | Coefficient | POPC | DPPC |
| --- | --- | --- | --- |
| Bond | $k_{H1-H2}$ | 0.446 | 0.471 |
| | $k_{H2-T1}$ | 1.073 | 1.320 |
| | $k_{T1-T2}$ | 1.001 | 0.875 |
| | $k_{T2-T3}$ | 0.443 | 0.280 |
| | $b_{0,H1-H2}$ | 5.580 | 5.417 |
| | $b_{0,H2-T1}$ | 5.452 | 5.824 |
| | $b_{0,T1-T2}$ | 5.050 | 6.312 |
| | $b_{0,T2-T3}$ | 5.095 | 6.299 |
| Angle | $k_{H1-H2-T1}$ | 0.600 | 0.582 |
| | $k_{H2-T1-T2}$ | 2.383 | 3.357 |
| | $k_{T1-T2-T3}$ | 0.880 | 4.823 |
| | $\theta_{0,H1-H2-T1}$ | 3.142 | 3.142 |
| | $\theta_{0,H2-T1-T2}$ | 3.142 | 3.142 |
| | $\theta_{0,T1-T2-T3}$ | 3.142 | 3.142 |

The two remaining terms of the force field represent the interaction between two lipid molecules. These terms have the same functional form as the ones described in the work of Cooke, Kremer, and Deserno^10^. The repulsive term, *V*_*Repulsion*_, is modeled with the Weeks-Chandler-Andersen potential:

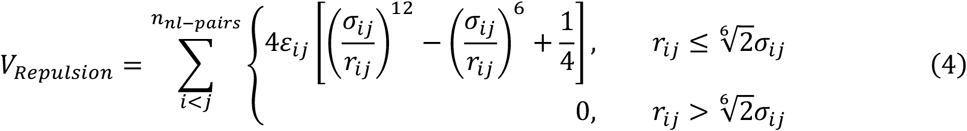

with *ε*_*ij*_ representing the force scaling factor, *σ*_*ij*_ the repulsive range for the *ij* pair of beads, and *n*_*nl*−*pairs*_ the total number of non-local pairs of CG beads. This repulsive interaction is applied to all the pairs of beads that are not participating in a virtual bond or a bond-angle interaction. The value of *σ*_*ij*_ is defined as the arithmetic mean (*σ*_*i*_ + *σ*_*j*_)/2 where *σ*_*i*_ (*σ*_*j*_) represents the van der Waals diameter of the *i*-th (*j*-th) CG particle. For each lipid type, *σ*_*i*_ takes two values; *σ*_*H*_ for the head and *σ*_*T*_ for the tail beads, with values related by *σ*_*H*_ = 0.65*σ*_*T*_ (**Fig. 1b**). This relation confers the lipids a geometry that prevents the formation of persistent holes in membranes, which we will discuss later in this paper. The value of *ε*_*ij*_ is defined as the geometric mean, 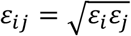, where *ε*_*i*_ depends on the lipid type. Finally, in the last term, *V*_*Attraction*_, represents the attractive hydrophobic interaction between tail beads of different lipid molecules:

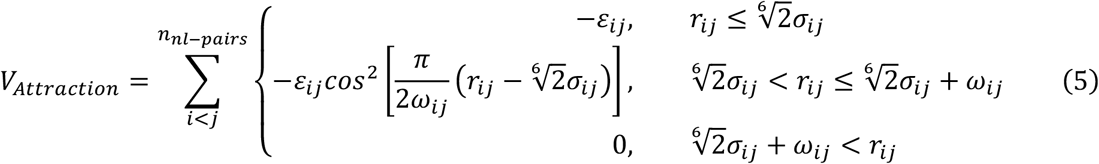

Here, *ε*_*ij*_ and *σ*_*ij*_ have the same values as in the repulsive potential, whereas *ω*_*ij*_ represents the width of the pair potential well, which is defined as the arithmetic mean of each lipid type, (*ω*_*i*_ + *ω*_*j*_)/2. The attractive interaction, which approximates the hydrophobic interaction, is only applied between tail bead pairs, excluding the T1-T3 pairs (**Fig. 1b**). This exclusion of the T1-T3 pairs was motivated to prevent the system from getting trapped into a highly rough membrane surface, and unrealistic collapsed local minima. **Fig. 1c** shows the potential that represents the interaction between tail bead pairs that participate in both repulsive and attractive interactions. In our preliminary tests, an attractive potential between head beads occasionally resulted in the inverted micelle form for simulations starting from random configurations. To avoid this configuration, we do not include the attraction between head beads in the current model. However, a weak and fine-tuned attractive force could be useful in the future.

We call the current CG lipid models as the *i*mplicit *so*lvent *l*ipid *f*orce field, iSoLF.

### Coarse-grained molecular dynamics simulation

We performed all the CG MD simulations using a modified version of our software, CafeMol v3.1.^38^ We used the standard underdamped Langevin equation and set the mass of each CG bead equal to the sum of the masses of atoms involved in the CG bead.

For the force-field parameter optimization, the estimate of physical properties of plane-membrane, their temperature dependence, the observation of pore formation, and the observation of the POPC/DPPC two-component membrane behavior, we used periodic boundary conditions and semi-isotropic pressure coupling in the xy-direction by fixing the linear length in the z-axis and allowing the linear lengths in the x- and y-axes to change while maintaining the surface tension equal to zero, i.e., the Nγ_xy_L_z_T ensemble where γ_xy_ means surface tension in xy-direction and Lz stands for the linear length in the z-direction. For the integrator, we used the one developed by Gao, Fang, and Wang.^39^ The friction coefficient of the thermostat in the Langevin dynamics was set equal to 0.1 (1/CafeMol-time. In CafeMol v3.1, one CafeMol-time unit apparently corresponds to ∼ 49 fs although effective dynamics is much accelerated.) ^38^, the friction coefficient of the barostat equal to 0.1 (1/CafeMol-time), and the compressibility of the simulation box equal to 0.01 (Å^3^ · *mol*/*kcal*). The MD time-step size in the integration was 0.2 (CafeMol-time) for simulations of pure POPC system, and 0.1 for simulations containing DPPC (we found the DPPC-containing system unstable with a time-step of 0.2, probably due to a small harmonic force constant and a large bond length of T2-T3 bonds). For the parameter optimization process, each simulation consists of 1 × 10^6^ and 2 × 10^6^ MD steps for POPC and DPPC, respectively, of which the second half data were used for the estimation of the properties. For the temperature dependence examination, each simulation contains 1.2 × 10^6^ and 2.4 × 10^6^ MD steps for POPC and DPPC systems, respectively, from which the first sixth of data was discarded (sample trajectories are in **Fig. S1**). For the POPC/DPPC two-component system, the simulation contains 2.4 × 10^6^ MD steps.

For the spontaneous lipid bilayer membrane formation and the vesicle simulation, we used the default dynamics setup of CafeMol, a fixed-size box with periodic boundary conditions, the NVT ensemble, and 2 × 10^6^ and 1 × 10^6^ MD steps, respectively.

For the simulation of the spontaneous lipid bilayer membrane formation, we prepared the initial configuration by sequentially placing lipid molecules inside a simulation box. The placement procedure of lipids consisted of selecting a random point where an H1 head bead was positioned. Then, the remaining beads (H2, T1, T2, and T3) were added following a randomly oriented straight line. During the placement of a lipid molecule, if any of its beads happened to be at a distance lower than 1.3*σ* to any of the already placed beads, the molecule was discarded, and the placement of the H1 bead was performed again.

For the simulation of a vesicle made of POPC lipids, we prepared an initial configuration based on a simple geometric method. We set the radius of the vesicle as 15 nm (more precisely, the radius is defined as the distance from the center to the outer layer of the vesicle). Using the surface area of each leaflet and the area per lipid, we estimated the number of lipids in the inner- and outer-leaflets as 2976 and 4400, respectively. Then, we used an algorithm that distributes points optimally on the surface of a sphere as evenly as possible, via mapping the Fibonacci lattice onto the surface^40^, by which we placed each lipid molecule.

We implemented our lipid model, iSoLF, in our software, CafeMol, and it will be included in the upcoming release.

### All-atom molecular dynamics simulations

For the bottom-up parameter determination and calculation of reference physical properties, we performed all-atom MD simulation using GROMACS^41^ version 5.1.1, the Slipids^42,43^ force field for lipids, and the TIP3P water model.^44^ We downloaded single-component membrane patches of 128 POPC and DPPC lipids (64 lipids per leaflet) from the Slipids website^45^ and performed energy minimization using the steepest descent method, NVT equilibration for 200 *ps* at a constant temperature of 303K using the v-rescale thermostat, and NPT equilibration for 5 *ns* at a constant pressure of 1.013 bar and a constant temperature of 303K using the Parrinello-Rahman barostat. In both equilibrations, lipid and water molecules were coupled separately, using time constants of 0.5ps and 10ps for the thermostat and barostat, respectively. Then, we performed production runs for 200 *ns* using the equilibrated patches, of which data was used for the estimates.

Finally, for the running-time comparison of the CG and all-atom models, we equilibrated all-atom and CG membrane patches of POPC following the protocol described above and performed production runs to estimate 2D MSD using 1 CPU core of an Intel i7-5930K processor and no GPUs.

### Calculations of properties

In the planar membrane simulations, we calculated the area per lipid, the order parameter, the hydrophobic thickness, and the 2D diffusion coefficient. The area per lipid, *A*_*L*_, was calculated by the formula:

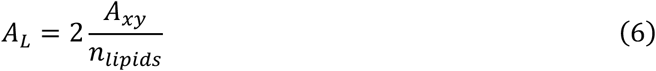

where *A*_*xy*_ is the area of the simulation box in the xy-plane and *n*_*lipids*_ is the total number of lipids in the system.

The order parameter, *S*_*θ*_, was calculated from the angle *θ*_*i*_ formed between the line joining the center of mass of the tail beads T1 and T3, and the z-axis (**Fig. 2a**). We used this angle in the formula:

**Figure 2.**
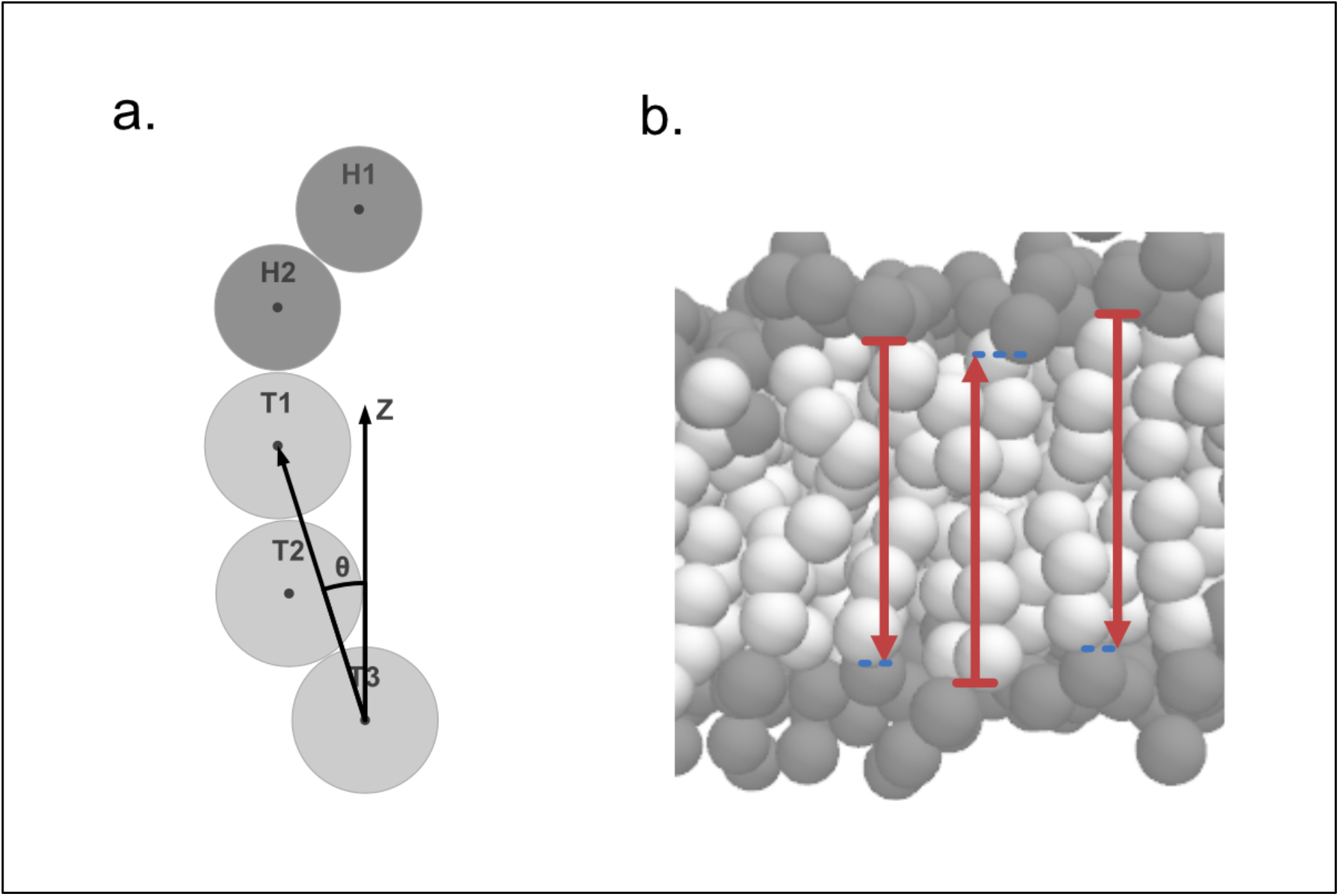
Methods for calculating the order parameter and the hydrophobic thickness. (a) Drawing of the angle *θ* for the calculation of the order parameter. It is defined as the angle between the line joining the bead T1 and T3, and the z-axis. (b) The local hydrophobic thickness for three lipids. For each lipid molecule, we find a lipid molecule in the opposite leaflet such that the distance in xy-plane between the two lipids is the smallest (blue dashed lines). The difference in the z-axis between the pair of lipids (indicated by red arrows) defines the hydrophobic thickness at that site.

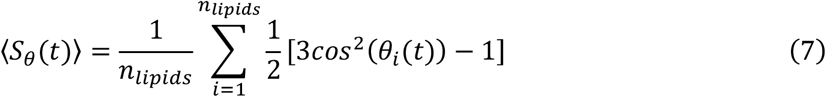

where *n*_*lipids*_ is the total number of lipids in the system, and ⟨⋯ ⟩ represents the average over all lipids.

For the hydrophobic thickness, we employed a similar method to the one used in the GridMAT-MD^46^ software. For each lipid molecule, we choose the middle point of the H2-T1 pair as a reference point. Then, we find the lipid molecule in the opposite leaflet, which has the smallest distance in the xy-plane. The difference in the z-axis between the reference points defines the hydrophobic thickness at that lipid site. Once it is averaged over all lipids, we obtain the hydrophobic thickness (**Fig. 2b**).

The 2D diffusion coefficient, i.e., the lateral diffusion coefficient, was calculated from the mean square displacement (MSD), defined as MSD = ⟨(*r*_*i,xy*_ (*t* + *t*_0_) − *r*_*i,xy*_ (*t*_0_))^2^⟩, where *r*_*i,xy*_ is the xy coordinate of the center of mass of the i-th lipid molecule, *t* is the size of the time window, and *t*_0_ is the separation between time windows. The MSD as a function of the time window *t* is fitted by a straight line of which the slope is equal to 4 times the 2D diffusion coefficient. Also, for a given *t*, we set the interval of *t*_0_ larger than *t*.^47^

Finally, in this study, we used the physical properties described above to assign a phase to each of the simulated lipid bilayers. The gel phase was characterized by markedly slower lateral diffusion of lipids and a high order parameter. Faster lateral diffusion of lipids corresponded to the liquid phase, in which the liquid disordered and the liquid ordered phases were characterized by a low and high order parameter, respectively. As a function of temperature, we monitored the lateral diffusion coefficient, the order parameter, as well as the hydrophobic thickness and the area per lipid. For each lipid membrane, we found significant changes in all these properties nearly around a temperature, which we identified as the phase transition temperature. The thresholds for the slower/faster lateral diffusion and the lower/higher order parameter are defined by the corresponding values at the phase transition temperature.

## Results and Discussion

### Model Parameterization

Parameters in the force field were determined for two target glycerophospholipids, POPC (1-palmitoyl-2-oleoyl-sn-glycero-3-phosphocholine) and DPPC (1,2-dipalmitoyl-sn-glycero-3-phosphatidylcholine). It has been characterized that near-physiological temperature (30 °C) single-component membranes of POPC and DPPC lipids exhibit liquid disordered and gel phases, respectively. Thus, these two lipids can serve as representatives of the gel and liquid disordered phases of lipid membranes. In the parameter determination, we used a partly bottom-up and partly top-down approach.

For the virtual-bond and bond-angle potential parameters, we took a bottom-up approach. We first performed all-atom simulations of single-component lipid membranes of POPC and DPPC (see the section “All-atom molecular dynamics simulations”). From the obtained structure ensembles, we fitted **Eq**. (2) and **Eq**. (3) using the standard Boltzmann inversion method.^48^ The obtained parameters are listed in **Table 1**. It should be noted that we solely used lipid structural samples in the bilayer membrane. Thus, we expect the obtained lipid parameters are appropriate for lipids in the membrane, but not necessarily for those out of membranes.

For the parameterization of the intermolecular repulsive and attractive potentials, we took a top-down approach, in which we iteratively optimized the parameters so that the CG MD simulations reproduced three, either physical or geometrical properties of lipid membranes. To guide this optimization process, we introduced a cost function:

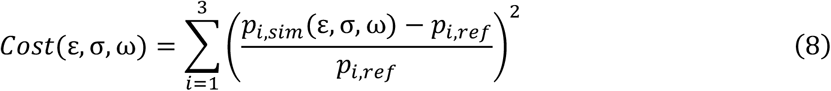

with *p*_*i*,ref_ representing the reference value for the i-th property and *p*_*i,si*M_ the i-th property calculated from the CG MD simulation that depends on the force field parameters, ε, σ, and ω. For the three properties to match, we chose the area per lipid (APL), the hydrophobic thickness, and the order parameter in the bilayer membrane (**Fig. 3a-3c**). The reference values for the first two properties for the POPC lipid membrane were taken from experimental data reported by Kučerka, Nieh, and Katsaras^49^. In contrast, those for DPPC in the gel phase that were not available from experiments were taken from the all-atom MD structure ensemble. We calculated the reference value for the order parameter from the all-atom MD trajectories by mapping the atomic coordinates of lipid molecules to the CG beads and using the **Eq**. (7).

**Figure 3.**
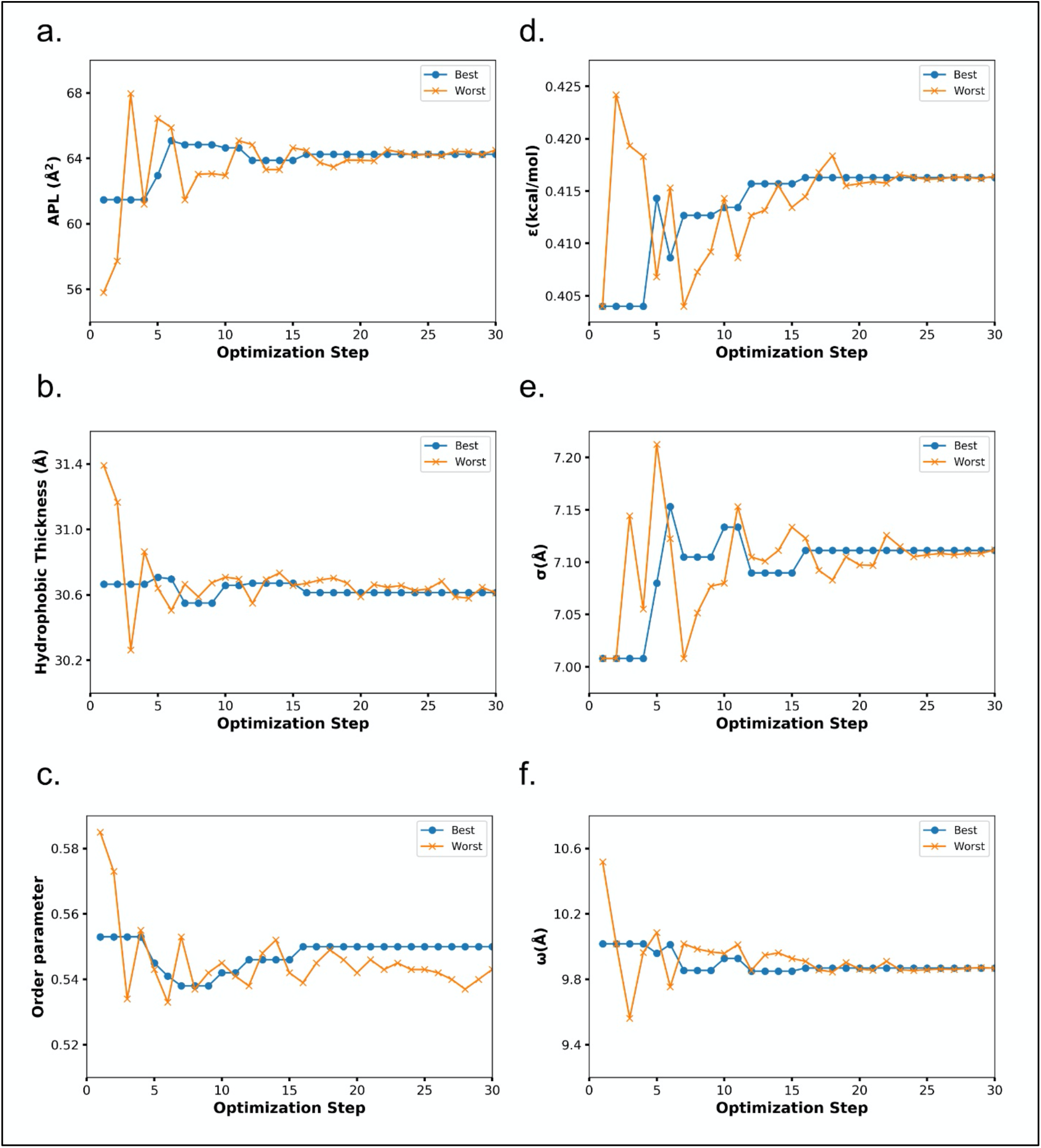
The parameter optimization process for the attractive and repulsive inter-lipid interactions for POPC using the Nelder-Mead method. (a)-(c) Values for the target properties for the best and the worst points of the boundary enclosing the minimum. As the optimization proceeds, both points approach each other. (d)-(f) Values for the coefficients of the attractive and repulsive interaction at each step of the optimization process.

We seek a set of parameters that minimize the cost function. Each evaluation of this cost function requires a new CG MD sampling with an updated set of parameters. Moreover, the calculation of the derivatives of the cost function with respect to the parameters is computationally very expensive. Thus, to minimize this function with respect to parameters, we used the Nelder-Mead method^50^, a gradient-free method in which a boundary enclosing a minimum in the parameter space is refined in each iteration step up to the desired precision. By selecting a suitable set of initial values, the convergence is achieved within some tenths of iterations (**Fig. 3d-3f**). Values for the optimized parameters for POPC and DPPC lipids are given in **Table 2**.

**Table 2.** Coefficients for intermolecular interactions of POPC and DPPC. *ε* is in kcal/mol, and *σ* and *ω* are in Å.

|  | POPC | DPPC |
| --- | --- | --- |
| $\epsilon$ | 0.416 | 0.464 |
| $\sigma_T$ | 7.111 | 6.900 |
| $\omega$ | 9.867 | 10.318 |

### Spontaneous membrane formation

Using the optimized set of parameters, we examined the spontaneous formation of lipid bilayer membranes with our CG force field, iSoLF. We prepared a system containing 200 POPC lipid molecules randomly placed in a box following the procedure described in the Methods section. We fixed the size of the box so it would produce the equilibrium APL of POPC at 30 °C, that is, L_x_, L_y_, and L_z_ equal to 64, 64, and 80 Å, respectively. With this setup, the randomly positioned lipids spontaneously adopted a lipid bilayer conformation within 10^4^ MD steps (**Fig. 4**). Once the lipid bilayer membrane was formed, we did not find any breakage of the membrane within the simulation time due to its stability. This spontaneous formation of the lipid bilayer membrane and no rupture of it suggest that the lipid bilayer is thermodynamically the most stable state with the current force field and under these conditions.

**Figure 4.**
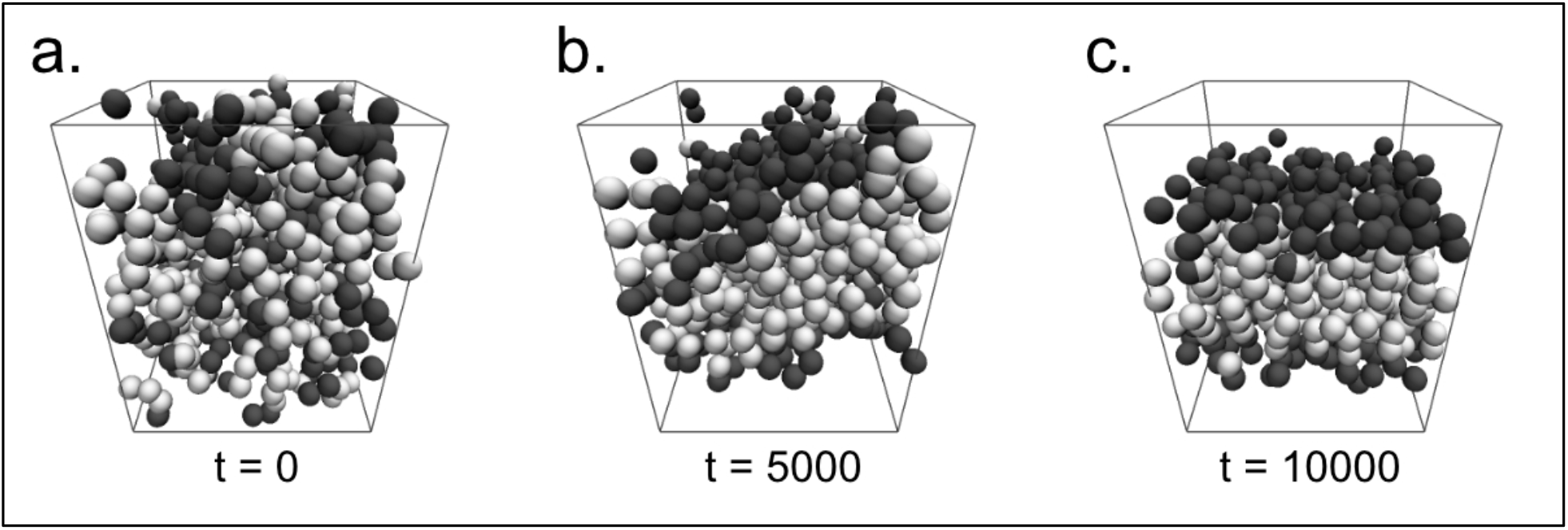
Spontaneous lipid bilayer membrane formation. (a) Simulation of 200 POPC lipids at 30 °C, starting from a random configuration. (b) After 5000 CafeMol-time units, lipids begin to gather, forming a membrane-like conformation. (c) The lipids adopt a membrane conformation after 10000 CafeMol-time units and maintain it without any breakage. Head and tail beads are in dark-gray and white, respectively.

Two observations in the preliminary studies may be instructive. First, when we used the ensemble of zero surface tension in the xy-direction with variable box size, we noticed that CG MD simulations starting from random conformations were unstable, and the system box expanded indefinitely. This might be a technical difficulty specific to the implicit solvent nature of a system with variable box size. To avoid this issue, we decided to use a fixed box size, i.e., the NVT ensemble, in this simulation.

Another interesting observation in the preliminary simulation is on the stability of the formed lipid bilayer membrane. With the choice of *σ*_*H*_ = 0.85*σ*_T_∼1.00*σ*_*T*_, during the simulations starting from the lipid bilayer membrane configurations, pores appeared spontaneously in the membrane (**Fig. 5a** and **Fig. S2**). This behavior is consistent with an observation given in a document by the original author group.^51^ When the box size is variable, the system occasionally expands, which results in a transient cavity formation in the membrane. The transiently formed cavity induces a tilt of the surrounding lipids, which increases the repulsive energy between head beads and tail beads at the cavity (akin to collisions). In order to reduce the repulsion, the system expands and forms a pore. Once the pore is formed in the membrane, it is very stable, and the system is trapped in this conformation (**Fig. 5a**). We found that the stability of the pores depends on the ratio *σ*_*H*_/*σ*_*T*_ between the head and tail beads (**Fig. S2**). With the choice of *σ*_*H*_ = 0.65*σ*_*T*_, we did not see, even transiently, the pore formation. Moreover, after a pore is formed with the use of *σ*_*H*_ = 0.85*σ*_*T*_ relation, by changing the ratio to *σ*_*H*_ = 0.65*σ*_*T*_, we could observe that the pore quickly disappeared (**Fig. 5b**). We concluded that a small enough radius of the head bead relative to the tail bead is necessary to make lipid bilayer membranes stable.

**Figure 5.**
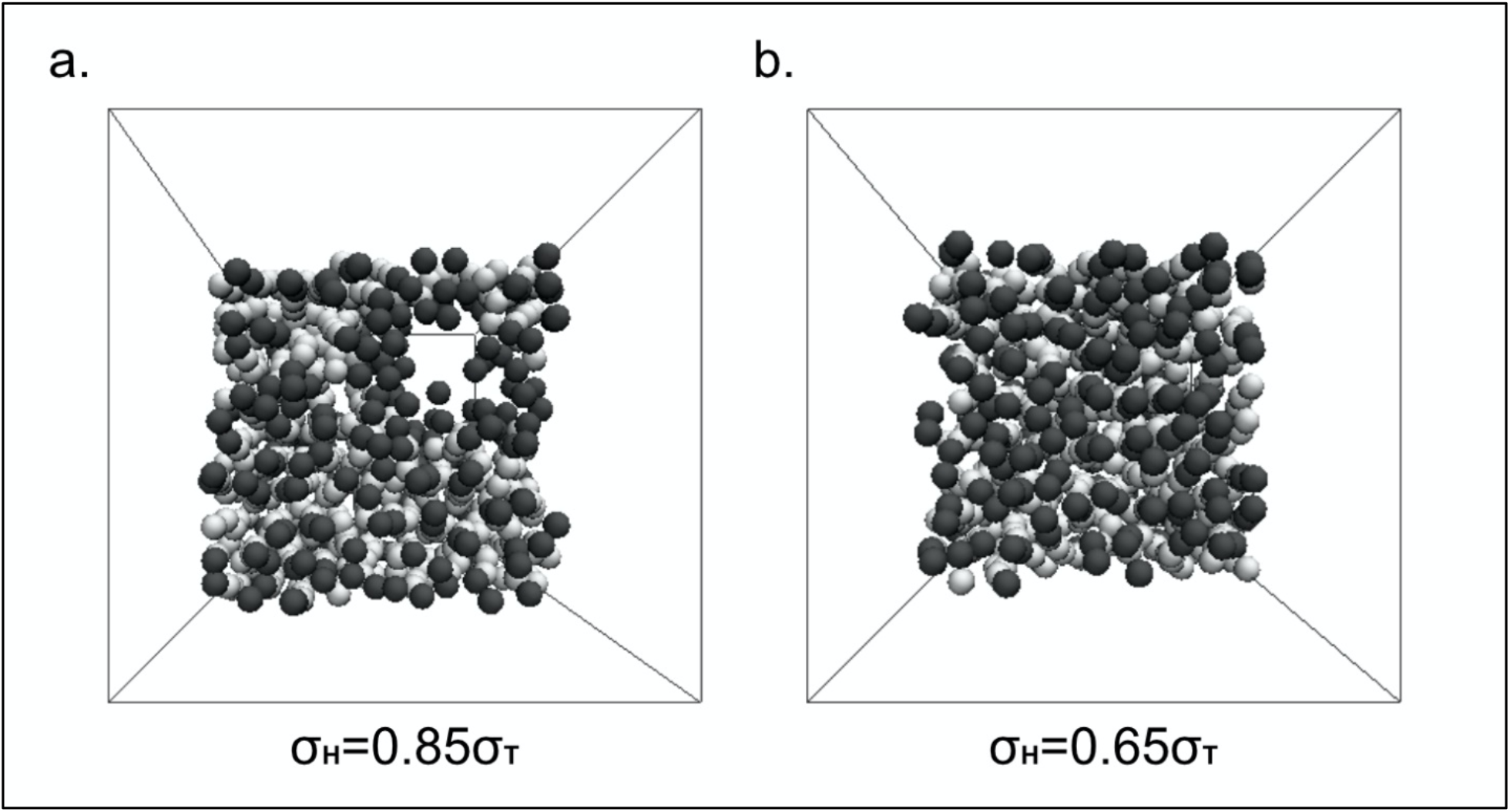
Pore formation in lipid membranes. Depending on the size ratio *σ*_*H*_/*σ*_*T*_, pores appear or disappear in the membrane conformations. (a) A simulation with a ratio of *σ*_*H*_ = 0.85*σ*_*T*_ results in the spontaneous formation of pores. (b) When the ratio is changed back to *σ*_*H*_ = 0.65*σ*_*T*_, the pore disappears. Head and tail beads are in dark-gray and white, respectively.

### Lateral diffusion

We evaluated the lateral diffusion of POPC and DPPC lipids with the parameter set determined in the optimization process. To quantify the lateral diffusion, we computed the MSD in 2D at 30 °C (**Fig. 6a**). The MSD with respect to the time difference fits well to the straight line, suggesting a normal diffusion in 2D. A comparison of the slope of the MSD of the two lipids suggests that the pure POPC membrane is in a liquid phase, whereas the pure DPPC membrane is in a gel phase at 30 °C. To further support this, we calculated the diffusion coefficient of POPC and DPPC at different temperatures (**Fig. 6b**) and observed an apparent phase transition from gel to liquid phases around 25 °C for POPC and 95 °C for DPPC. This putative assignment of phases is further confirmed later by observing simultaneous changes in the area per lipid, the hydrophobic thickness, and the order parameter.

**Figure 6.**
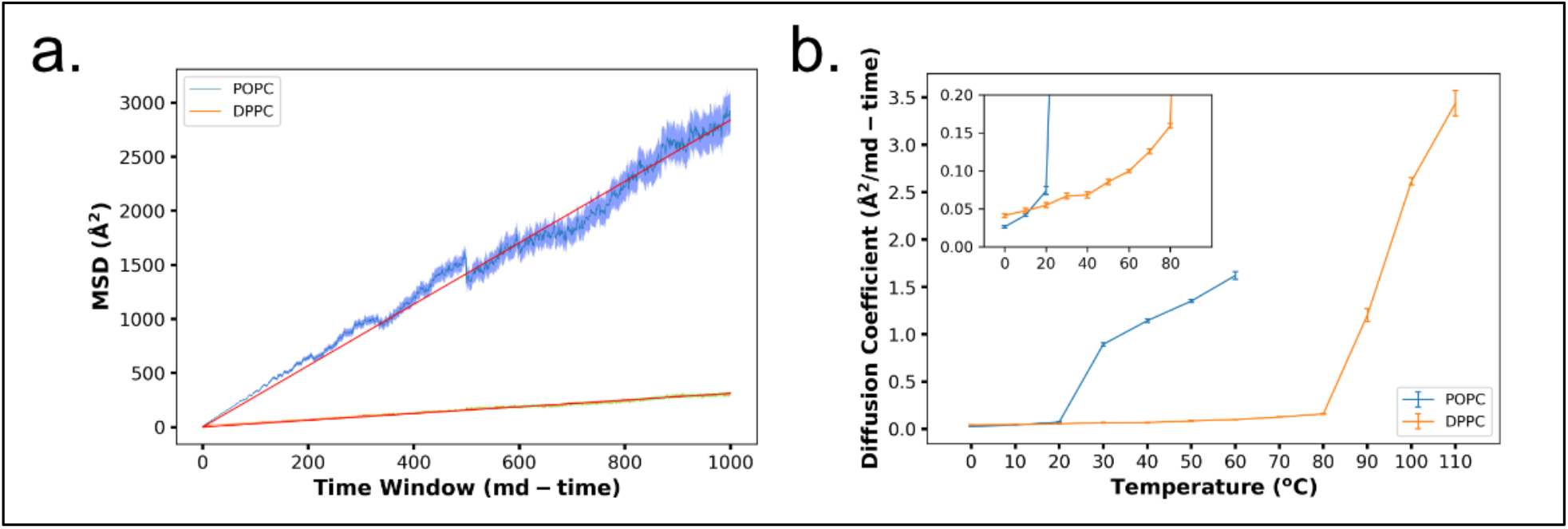
Lateral diffusion of POPC and DPPC lipids in membranes. (a) Mean square displacement (MSD) at a temperature of 30 °C for POPC (blue) and DPPC (orange). At this temperature, the POPC membrane shows a liquid phase, while the DPPC membrane remains in a gel phase. The red line corresponds to the fitted equation used to calculate the diffusion coefficient. (b) The 2D diffusion coefficient for POPC and DPPC lipids as a function of temperature. The model POPC and DPPC membranes exhibited an apparent phase transition around 25 °C and 95 °C, respectively. The subplot shows the lower section of the main plot.

We also used the MSD in 2D for a rough comparison of the running time of our coarse-grained lipid model with a standard all-atom model. Using 1 CPU core for both all-atom and coarse-grained simulations, we calculated the MSDs (**Fig. S3**). For the all-atom simulation, an MSD of 0.174 *nm*^2^ was obtained in about 8 *hours* 47 *minutes*. On the other hand, with our CG model, an MSD of 36.7 *nm*^2^ was obtained in about 21 *minutes*. Assuming the MSD increases linearly in time, except for a very short time regime, we obtained a speed-up factor of ∼5000 relative to the all-atom model. To get an MSD value comparable to our coarse-grained model, the all-atom model would need about 2 months and 17 days using the same resources.

### Vesicle dynamics

Next, we performed a CG MD simulation of a vesicle made of POPC lipids. For this, we prepared a small unilamellar vesicle (SUV) with a diameter of ∼30 nm (see the Method, **Fig. 7a**). Preliminary tests suggested that starting the CG MD simulations at a room temperature causes an unstable behavior of the vesicle due perhaps to a poor setup of the initial structure. To avoid this instability, we started from a temperature of 0K = −273 °C and then gradually heated the system until 30 °C. During this heating process, some lipid molecules in the outer leaflet left the vesicle, but without affecting the overall vesicle shape. Once the system reached 30 °C (**Fig. 7b**), we removed lipid molecules that were dissociated from the vesicle during the heating up process and, after that, conducted the production run. During this process, we evaluated the stability of the vesicle by monitoring the radius of the sphere that best fitted the vesicle as a function of time (**Fig. 7c**). Also, by visual inspection, we checked that the vesicle did not present any pore and that the unilamellar structure was maintained during the simulation.

**Figure 7.**
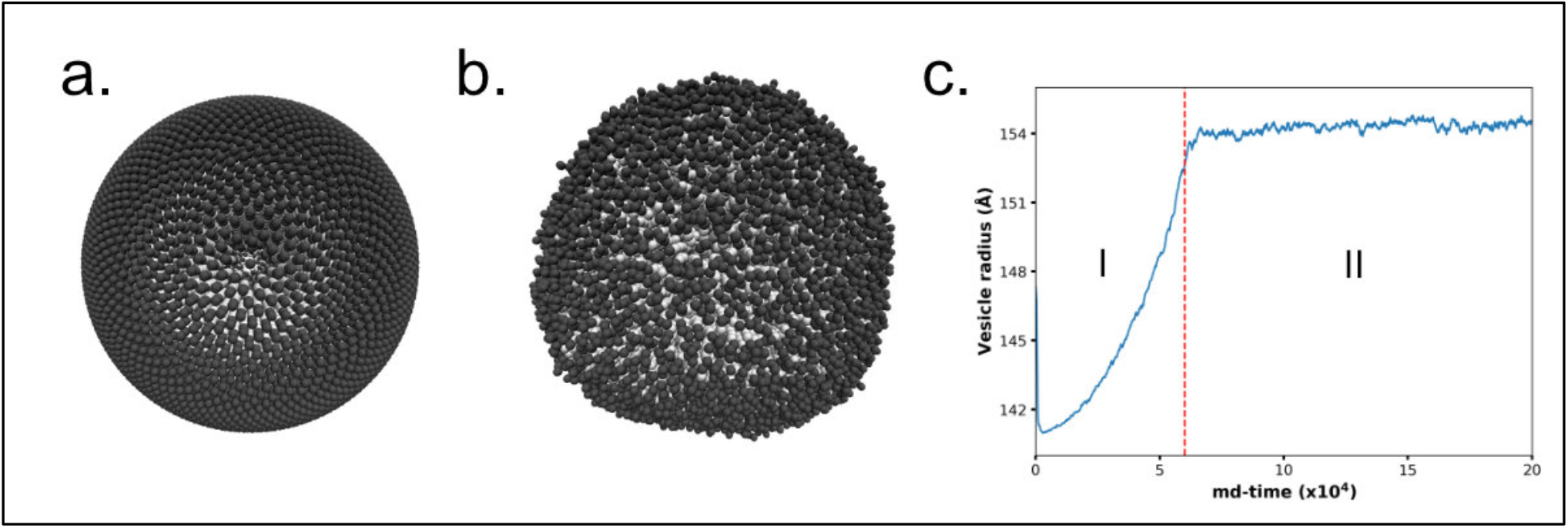
A CG MD simulation of a vesicle made of POPC lipids. (a) Initial conformation of the vesicle generated with the Fibonacci lattice method. (b) Vesicle after equilibrating the system at room temperature. (c) The radius of the best fit sphere on the vesicle during the equilibration (I) and production (II) run. In the first stage, the vesicle was heated up from 0 K = −273 °C to 30 °C, and in the second stage, the vesicle was kept at a constant temperature of 30 °C. Head and tail beads are in dark-gray and white, respectively.

### Temperature dependence

The parameterization of our CG lipid force fields for POPC and DPPC was performed at 30 °C. At this temperature, the simulations closely reproduced the reference properties, and the POPC and DPPC membranes were apparently in liquid and gel phases, respectively (**Fig. 3** and **Fig. 6b**). Here, we examine if our model can reproduce the temperature dependence of these quantities. Simulating POPC and DPPC lipid membranes at different temperatures from 0 °C to 110 °C, we calculated the area per lipid (APL), the hydrophobic thickness, the order parameter (**Fig. 8**), and the lateral diffusion coefficient (**Fig. 6b**) of single-component lipid bilayers.

**Figure 8.**
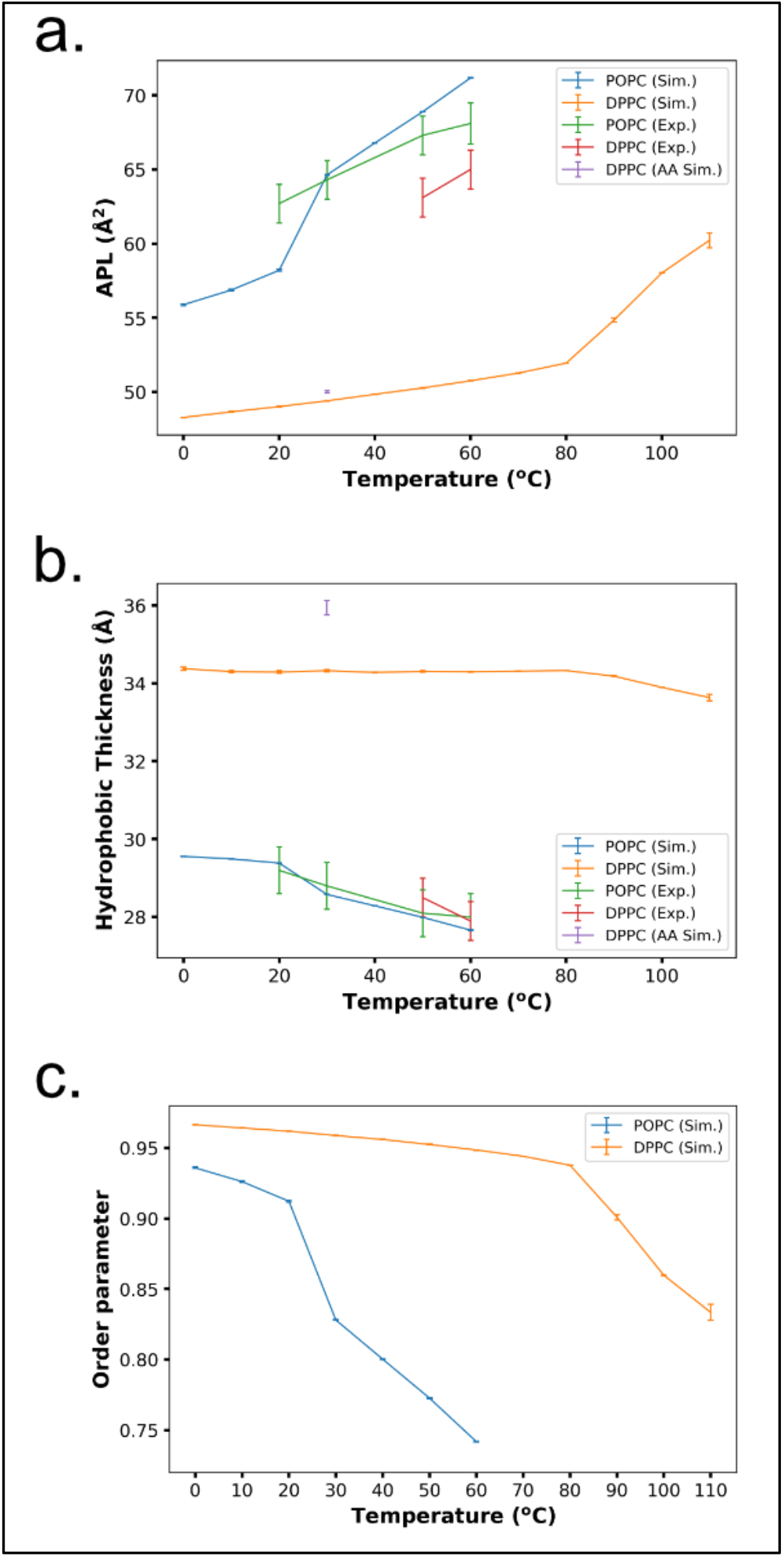
Temperature dependence of membrane properties. Comparison of the temperature dependence of geometric properties between CG simulations and experiments for (a) the area per lipid (APL), (b) the hydrophobic thickness, (c) and the order parameter. The experimental data are available only in the liquid phase for POPC. Purple squares represent the values calculated from all-atom MD simulations of DPPC membranes. The maximum errors are ±0.04 and ±0.009 for the APL, ±0.01 and ±0.004 for the hydrophobic thickness, and ±7.1 × 10^−4^ and ±0.006 for the order parameter of POPC and DPPC, respectively.

We observed a characteristic change in the area per lipid, the hydrophobic thickness, and the order parameter nearly at the same temperature as that in the lateral diffusion coefficient, both for POPC and DPPC (**Fig. 8**), giving further evidence of a transition from the gel phase to the liquid disordered phase^52-54^. For the pure POPC membrane, we found that both the area per lipid and the hydrophobic thickness of the CG model membrane stayed correlated with the experimental values at temperatures within the range of 30-60 °C. Below 30 °C, however, we observed the phase transition around 25 °C in the CG model (**Fig. 3** and **Fig. 6b**), whereas, experimentally, the phase transition temperature of the POPC membrane is reported as −2 °C. This shows that the transition temperature was not correctly reproduced in the current parametrization.

DPPC lipid membranes have a gel-liquid phase transition temperature at 41 °C, experimentally. In the current parametrization, we took the reference values at 30 °C from an all-atom MD simulation ensemble, in which DPPC lipid was in the gel phase. With our CG lipid force field, the DPPC lipid membrane showed a sharp gel-liquid phase transition at around 95 °C (**Fig. 6b**). Thus, the phase transition temperature was also not accurately reproduced. At 50-60 °C, our estimates of the area per lipid and the hydrophobic thickness from CG MD in the gel phase deviates from experimental data from the liquid phase (**Fig. 8**).

Overall, we summarize that, with the current force field of lipids, we can reproduce major geometrical properties of pure POPC and pure DPPC membranes at 30°C at which we calibrated the parameters, as well as the temperature dependence within the liquid phase. Yet, we cannot predict the phase transition temperature between the gel and liquid disordered phases.

### Two-component membrane system

Finally, we tested the behavior of a membrane composed of POPC and DPPC. We simulated a membrane consisting of 256 POPCs and 256 DPPCs at 50 °C. We prepared the initial configuration at which the POPC lipid molecules were localized in the half area of the membrane, and the DPPC lipids were localized in the other half (**Fig. 9a**). In an early stage of the simulation, we confirmed that two different phases, the liquid phase in the POPC region and the gel phases in the DPPC region, coexist (**Fig. 10a**).

**Figure 9.**
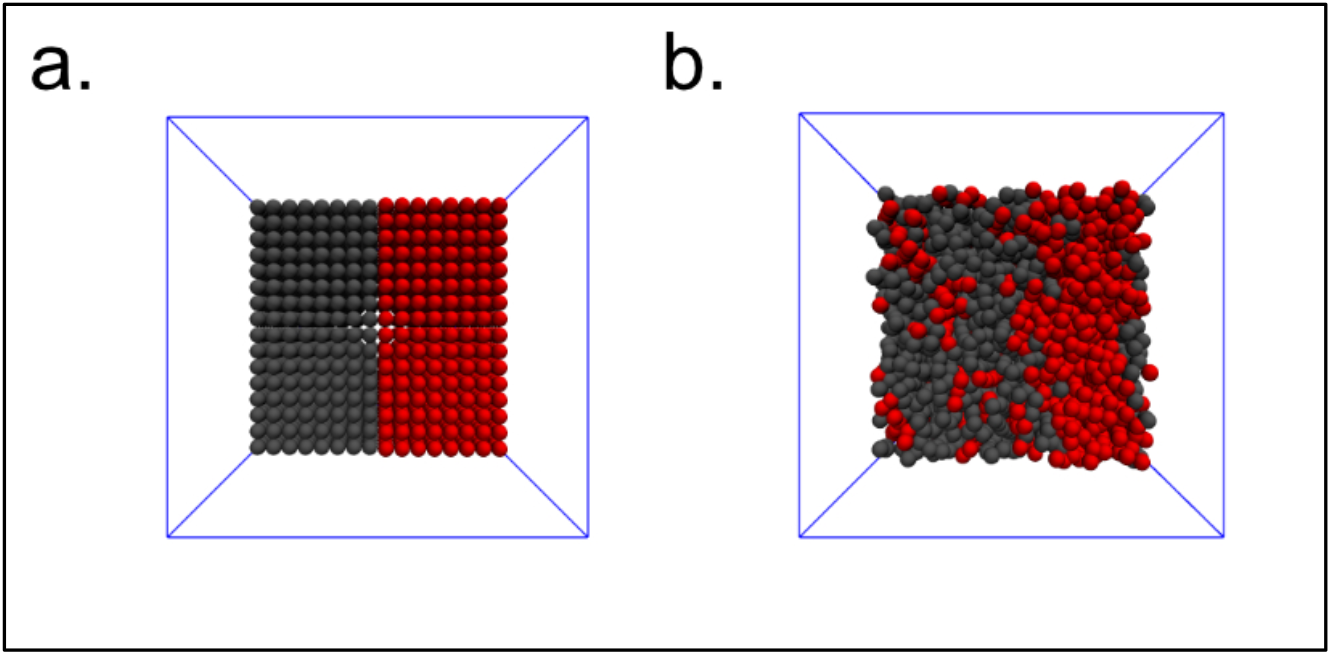
Simulation of a system composed of POPC and DPPC with a ratio of 1:1 at 30 °C. (a) Initial configuration of the binary system. Grey and red molecules represent POPC and DPPC lipids, respectively. (b) Some DPPC lipids in a gel phase diffuse to the POPC phase. At the boundary, POPC lipids directly interacting with DPPC lipids transitioned to a gel phase.

**Figure 10.**
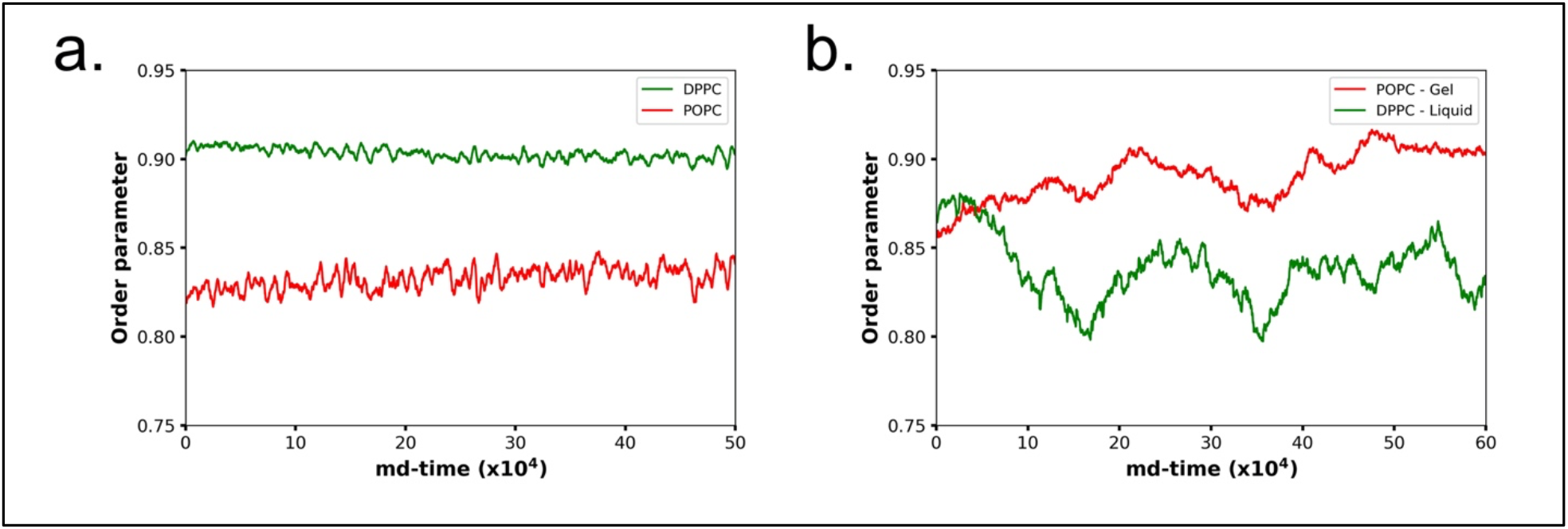
Order parameter for POPC and DPPC in a two-component membrane. (a) Time series of the average order parameter for POPC and DPPC in a two-component membrane. (b) Time series of the order parameter of two specific lipids. The red line shows the order parameter of a POPC lipid that diffuses into the DPPC gel phase, and the green line the order parameter of a DPPC lipid that diffuses into the POPC liquid phase.

As the system evolved, we observed some exchanges in lipids around the interface between the DPPC and POPC phases. We found that the DPPC lipids that moved to the liquid disordered phase exhibited a faster diffusion, whereas the POPC lipids that moved to the gel phase had almost zero diffusion. Consistently, the order parameter suggests that DPPC lipids exhibit liquid disordered-like behavior when they are locally in a low ratio to POPC lipids, despite being at a temperature that corresponds to the gel phase of pure DPPC (**Fig. 10b**). In the same way, POPC lipids exhibited a gel-like behavior when they are locally in a lower ratio to DPPC lipids.

## Conclusions

In this study, we extended the three-bead lipid model developed by Cooke, Kremer, and Deserno into a five-bead model. We parametrized it for the two phospholipids, one unsaturated, POPC, and the other saturated, DPPC lipids. The developed model, iSoLF, reproduced the area per lipid, the hydrophobic thickness, and the phase behaviors of the target phospholipids at 30 °C. Also, the model membranes of POPC and DPPC were in liquid disordered and gel phases, respectively, in accordance with experiments. We further examined the spontaneous formation of a lipid bilayer, the temperature dependence of physical properties, the vesicle dynamics, and the POPC/DPPC two-component membrane dynamics using the parameterized CG lipid model.

While our CG model membranes, both for POPC and DPPC lipids, reproduced geometric and physical properties estimated from experiments or all-atom models at 30 °C where we calibrated the parameters, the CG model did not reproduce the gel-liquid phase transition temperature correctly. Probably, we can perform finer tuning of the parameters targeting the phase transition temperature for each target phospholipid. For the two-component systems made of POPC and DPPC, we only tested a small patch of a membrane with 1:1 composition in this work. Probably, we need a more comprehensive examination of longer-time simulations of larger systems with different compositions. These refinements are left for future studies.

Since our aim here is to develop a CG lipid model compatible with the Cα-protein model, our next step is to model lipid-protein interactions. Therein, from a physicochemical point of view, the excluded volume, hydrophobic interactions, and electrostatic interactions need to be modeled. These developments are now underway. Once combined with standard Cα protein models, the iSoLF model will be a powerful tool to simulate large biological membrane systems made of lipids and proteins. The iSoLF model will be available in the upcoming release of CafeMol.

## Data Availability

The data that support the findings of this study are available from the corresponding author upon reasonable request.

## Author’s Contribution

All authors contributed equally to this work.

## Acknowledgment

We thank Hiroshi Noguchi for many helpful discussions, Giovanni Brandani, for his valuable insights, and Toru Niina, Suguru Kato, and Masahiro Shimizu for their help when implementing the model in CafeMol. D.U.L.T. acknowledges the Japanese Government Scholarship from the Ministry of Education, Culture, Sports, Science, and Technology (MEXT). The study was supported partly by MEXT as “Priority Issue on Post-K computer” (ST), by the RIKEN Pioneering Project “Dynamical Structural Biology” (ST) and by the Japan Science and Technology Agency (JST) grant JPMJCR1762 (ST).

## Supporting Information

**Figure S1.**
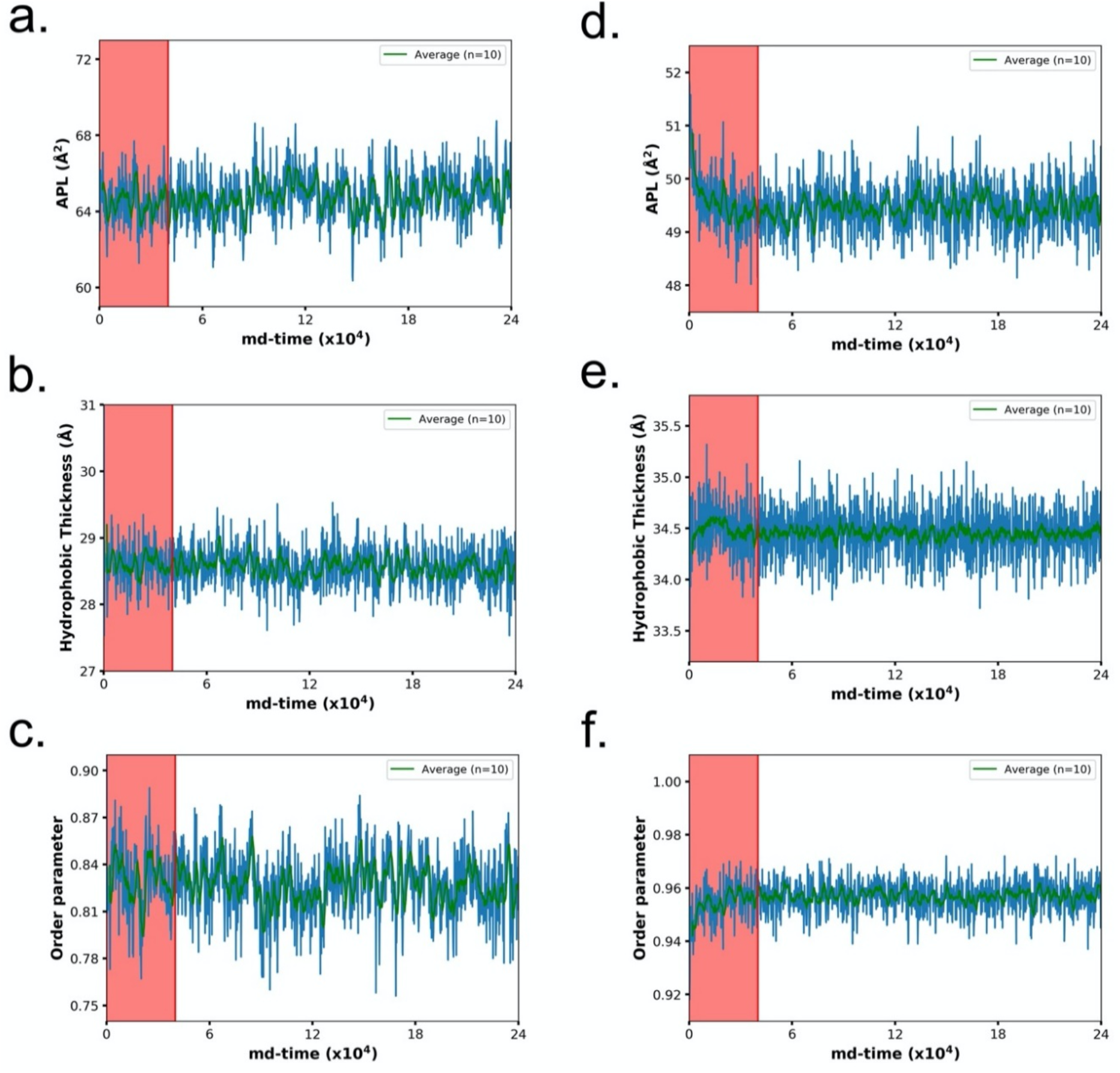
Time series for POPC and DPPC. Time series obtained with our coarse-grained lipid model following the protocol described in the Methods section of the main text. The plots show the APL, Hydrophobic Thickness, and Order parameter for POPC (**a-c**) and DPPC (**d-f**). The red zone in each plot represents the portion of the trajectory that was discarded, and the green line represents the running average for the 10 last points.

**Figure S2.**
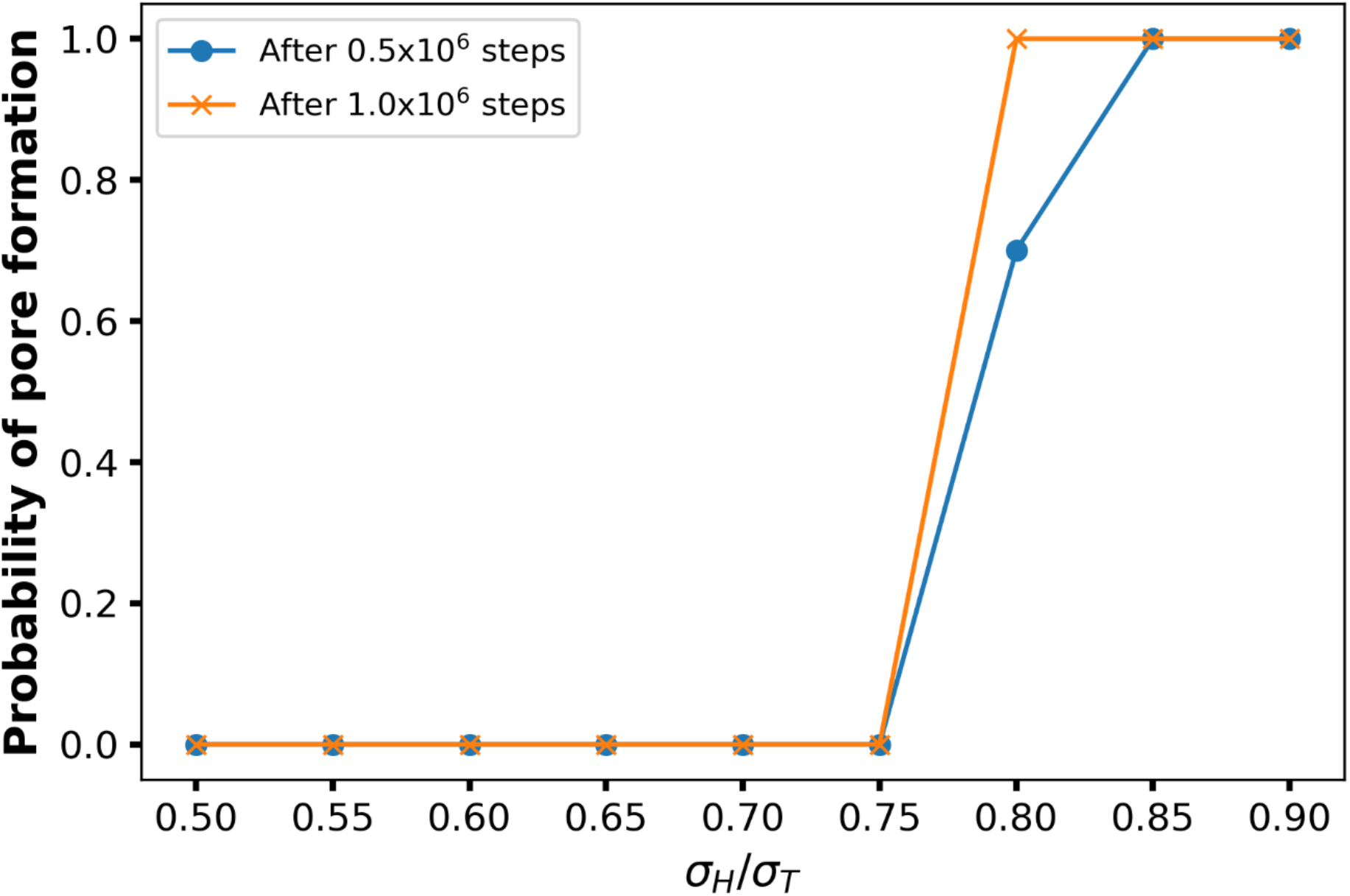
Probability of pore formation for POPC. For each ratio, 20 simulations were performed from which the probability was determined by counting the number of membranes presenting a pore. No pores were formed when the ratio of *σ*_*H*_/*σ*_*T*_ is 0.75 or lower. At a ratio of 0.8, a pore was formed in 14 out of 20 membranes during the first 0.5 × 10^6^ simulation steps (blue line), and after 1.0 × 10^6^ simulations steps, all the membranes presented a pore.

**Figure S3.**
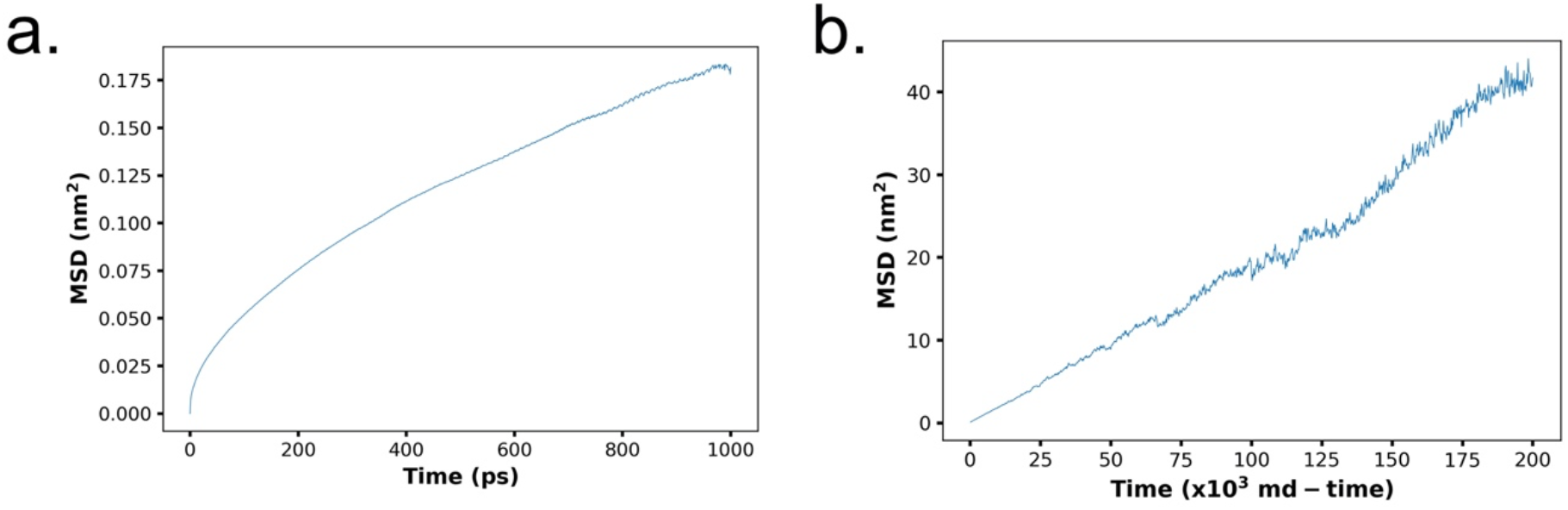
Lateral diffusion of POPC for one sample trajectory. (**a**) MSD obtained with the Slipids all-atom model. (**b**) MSD obtained with our iSoLF coarse-grained model. Both simulations consisted of 128 POPC lipids at 303K, as described in the Methods section of the main text.

