## Supporting Information for "Coarse-grained implicit solvent lipid force field with a compatible resolution to the Cα protein representation"

Diego Ugarte La Torre[1] and Shoji Takada[1]
[1]Department of Biophysics, Graduate School of Science, Kyoto University, Kyoto, Japan

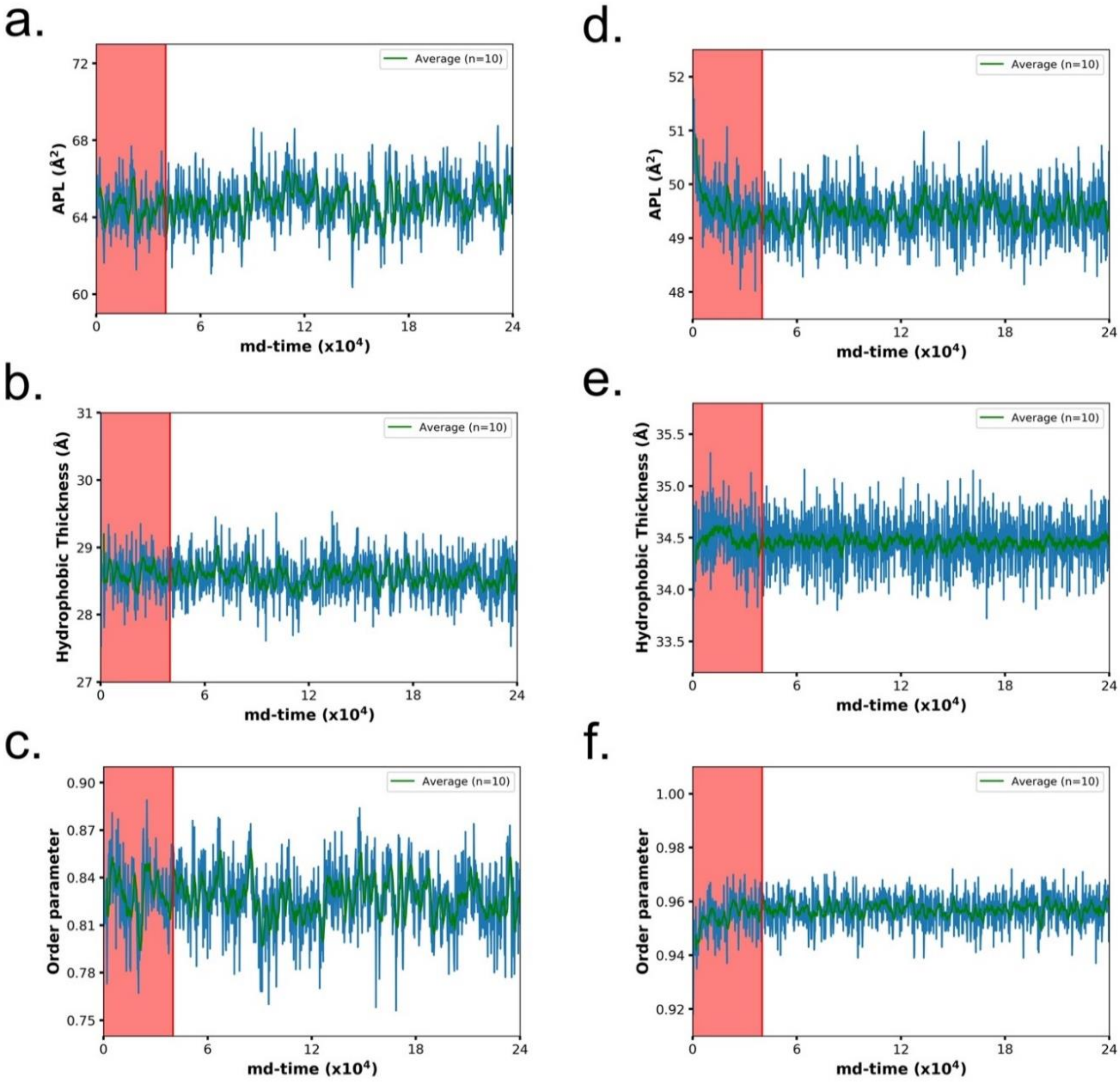


**Figure S1**. Time series for POPC and DPPC. Time series obtained with our coarse-grained lipid model following the protocol described in the Methods section of the main text. The plots show the APL, Hydrophobic Thickness, and Order parameter for POPC (**a-c**) and DPPC (**d-f**). The red zone in each plot represents the portion of the trajectory that was discarded, and the green line represents the running average for the 10 last points.

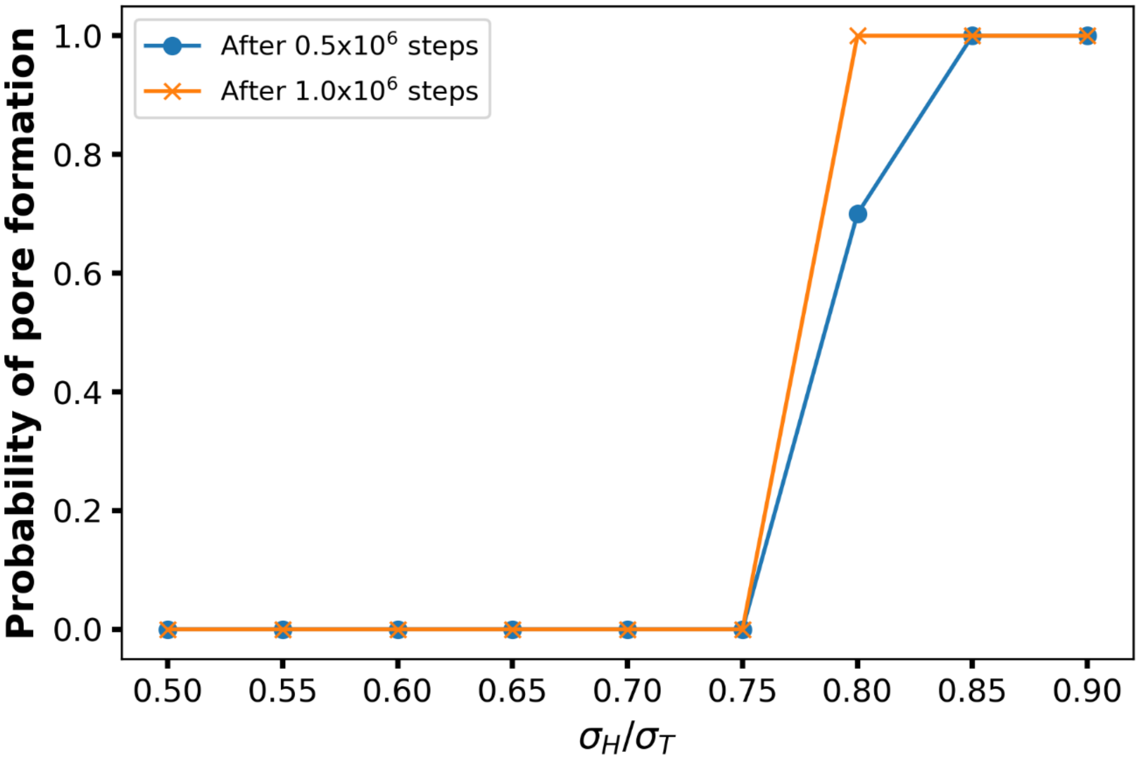


**Figure S2**. Probability of pore formation for POPC. For each ratio, 20 simulations were performed from which the probability was determined by counting the number of membranes presenting a pore. No pores were formed when the ratio of $\sigma_H/\sigma_T$ is 0.75 or lower. At a ratio of 0.8, a pore was formed in 14 out of 20 membranes during the first $0.5 \times 10^6$ simulation steps (blue line), and after $1.0 \times 10^6$ simulations steps, all the membranes presented a pore.

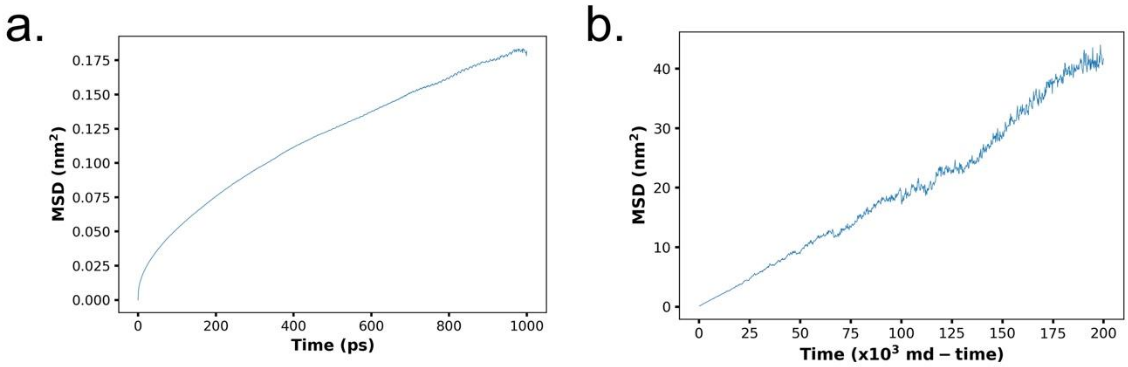


**Figure S3**. Lateral diffusion of POPC for one sample trajectory. (**a**) MSD obtained with the Slipids all-atom model. (**b**) MSD obtained with our iSoLF coarse-grained model. Both simulations consisted of 128 POPC lipids at 303K, as described in the Methods section of the main text.
